# Citizen Science Reveals Lepidopteran-Visitor Flower Network in Metro Manila, Philippines: A Preliminary Study

**DOI:** 10.64898/2026.02.17.706497

**Authors:** Samuel C. Brillo, Rodelina C. Deyto

## Abstract

Urban environments often harbor diverse pollinator assemblages, yet Lepidopteran-mediated pollination in tropical cities remains poorly documented. This preliminary study utilized research-grade iNaturalist observations (2015–2025) to construct a Lepidopteran visitor–flower network in Metro Manila, Philippines. A total of 126 validated records captured 38 lepidopteran taxa and 49 flowering plant taxa, predominantly exotic ornamentals and weedy species. Bipartite network analysis revealed a generalized interaction structure, with several butterfly species (*Papilio demoleus*, *Leptosia nina*, *Ypthima stellera*) and plants (*Tridax procumbens*, *Lantana camara*, *Ixora* spp.) serving as central hubs. Flower color–visitor heatmap indicated strong preferences for white, yellow, and pink flowers, while rare taxa demonstrated sporadic, low-frequency interactions. Notably, new nectar associations were documented for the endemic butterflies *Pareronia boebera*, *Troides rhadamantus*, and *Ypthima stellera*, highlighting the ecological role of cultivated and ruderal plants in sustaining endemic Lepidoptera within urban habitats. While opportunistic citizen science data are subject to spatial, temporal, and observer biases, they provide a cost-effective means of uncovering otherwise undocumented interactions, generating baseline insights, and informing urban pollinator conservation. These findings demonstrate the potential of integrating citizen science with ecological network analysis to characterize Lepidopteran–plant interactions and guide biodiversity-friendly urban planning in the city.

## INTRODUCTION

Pollination is a fundamental ecological process that sustains biodiversity, maintains plant reproductive success, and supports ecosystem resilience (Patil et al., 2024). While the majority of pollination studies have focused on bees as the dominant pollinators, other insect groups, such as Lepidoptera (butterflies and moths), also play significant yet often underappreciated roles in facilitating pollen transfer and plant reproduction (Chatterjee et al., 2024). Lepidopterans exhibit diverse behaviors and floral preferences that contribute to pollination across a wide range of angiosperm taxa, particularly in tropical and subtropical regions where floral diversity is exceptionally high (Dar et al., 2017). Despite this, Lepidopteran-mediated pollination remains poorly documented compared to bee–flower interactions, especially in urban landscapes undergoing rapid habitat modification (Ramírez-Restrepo & MacGregor-Fors, 2017).

Urbanization is one of the most pervasive drivers of biodiversity loss, leading to habitat fragmentation, reduction of native flora, and disruption of ecological interactions (Hald-Mortensen, 2023). In cities like Metro Manila, one of the most densely populated urban centers in Southeast Asia, green spaces are often limited to parks, gardens, and roadside vegetation, which is often dominated by ornamental and exotic species. Yet, these urban floral resources may still sustain diverse pollinator assemblages, including butterflies that have adapted to human-modified environments (López-Uribe et al., 2025). Understanding how Lepidopteran visitors interact with available floral resources in urban areas is essential for designing pollinator-friendly landscapes and conserving ecological processes within cities. Traditional field-based methods for studying pollination networks usually require extensive and systematic observation, which can be difficult to implement in complex urban environments. In this context, citizen science platforms such as iNaturalist offer a powerful alternative for documenting species interactions through opportunistic photographic observations (Bloom & Crowder, 2020). In addition, Citizen science plays an important role in identifying and monitoring pollinators, producing valuable information that supports their conservation and enhances our understanding of plant–pollinator relationships (Bitonto et al., 2025)

For instance, Beccacece et al. (2025) showed that diurnal arctiid moths primarily visited native Asteraceae species with rotate floral architecture and white-colored flowers, using at least 500 image-based observations. On the other hand, citizen science datasets used by Pernat et al., (2024) revealed that *Isodontia mexicana* exhibits distinct plant family preferences, with Lamiaceae—particularly *Pycnanthemum*—being the most frequently visited in North America. Analysis of flower colors from iNaturalist images further shows that this species predominantly visits white and yellow flowers across both North America and Europe. Fonturbel et al. (2023) investigated two exotic and one native species of bumblebees (*Bombus*) using data from citizen science. While ornamental exotic plants may act as foundations that facilitate their invasion, the exotic bumblebees mainly visited exotic plants and displayed preferences for particular flower colors. These recent studies reveal promising insights into pollinator biology by utilizing photographic observations from citizen scientists. Each image uploaded to iNaturalist represents a verifiable record of a biological encounter, and when such records capture Lepidopterans visiting flowers, they provide valuable opportunistic data for inferring potential plant–pollinator associations. Although such datasets are inherently non-systematic and subject to sampling bias, they can reveal baseline patterns of interaction and serve as a foundation for hypothesis generation and future field validation (Brown & Williams, 2019).

Our study aims to construct and describe a preliminary Lepidopteran visitor–flower network in Metro Manila derived from limited yet unique iNaturalist observations. Specifically, it seeks to (1) identify the Lepidopteran and plant species involved in observed visitation events, (2) characterize the structure and properties of the resulting interaction network, and (3) determine the origin status (native or exotic) of the visited plant species and the endemism status of the Lepidopteran visitors. The findings are expected to provide a baseline understanding of urban pollination patterns in Metro Manila and demonstrate the utility of citizen science data in documenting ecological interactions in data-deficient tropical cities.

## METHODOLOGY

### Data Collection

This study employed a citizen science–based research design, utilizing publicly available *iNaturalist* (https://www.inaturalist.org/) records as the primary data source. Because such data are opportunistic in nature, observations were contributed voluntarily by users rather than collected through standardized sampling protocols. This approach assumes that publicly shared photographic records reasonably represent real species interactions within accessible and urbanized environments. However, it may introduce potential sampling biases, such as uneven spatial coverage, observer preference toward conspicuous species, or underrepresentation of nocturnal taxa.

Despite these limitations, citizen science datasets offer a valuable and cost-effective resource for ecological research. They enable the discovery of otherwise undocumented species interactions and help fill knowledge gaps in areas where time, logistics, or accessibility constrain systematic field sampling. Thus, *iNaturalist* data are suitable for establishing preliminary insights into lepidopteran–plant associations within the urban landscapes of Metropolitan Manila.

Research-grade (RG) observations were obtained from an ongoing Community Project called “Lepidoptera of Metro Manila”, which can be found at this link: https://www.inaturalist.org/projects/lepidoptera-of-metro-manila?tab=observations on November 6, 2025, to document lepidopteran visitors associated with flowering plants within the area. Only observations meeting the following inclusion criteria were considered: (1) the observation must show a lepidopteran individual in direct contact with or landing on a flower; (2) the flower should visibly display reproductive structures or a position suggestive of a pollination interaction; and (3) the observation must be geographically recorded within the political boundaries of Metro Manila. Observations that did not meet these criteria were excluded from this study.

For each qualified research-grade observation, the following data were recorded: observer username, city (political boundary), month and year, time of day, lepidopteran visitor species, common name, lepidopteran family, associated plant species, plant family, and the prominent flower color (as seen in the corolla or in bracts functioning as attractive structures, e.g., *Mussaenda*, *Musa*, *Bougainvillea*). The endemism status of lepidopteran visitors was determined using available checklists, while the origin of the visited plant species (native or exotic) was verified based on global data from *Plants of the World Online* (POWO) and local data from *Co’s Digital Flora of the Philippines*.

### Data Cleaning and Verification

All records were manually inspected to ensure data quality and accuracy. Duplicate observations, misidentified taxa, or entries with unclear photographic evidence of flower visitation were removed. Taxonomic names of both lepidopteran visitors and plants were cross-checked and standardized following POWO and CoL (*Catalogue of Life*) to maintain consistency in nomenclature. When inconsistencies in identification occurred, higher taxonomic levels (e.g., family or genus) were retained to preserve valid interaction data.

When plant images closely resembled a particular species but could not be verified with full certainty, the identification was annotated with “cf.” to indicate a confident placement at the genus level while acknowledging uncertainty at the species level. Some RG observations showed unmatching metadata for time (e.g., the image depicts daytime but the metadata indicates nighttime); these were retained but marked as “N/A” for the time variable. The same treatment was applied to records with obscured or missing time stamps. For RG observations with obscured location data but still confirmed to fall within the boundaries of Metropolitan Manila, the closest or most precise location suggested by the iNaturalist distribution map was recorded.

### Data Analysis

The compiled dataset was analyzed using Python. A bipartite network was constructed to visualize the associations between lepidopteran visitor species and their recorded plant species, representing the interaction structure of the urban pollination network. Additionally, a heatmap was generated to illustrate the association between visitor species and flower color. These visualizations were used to identify potential interaction patterns and highlight commonly visited plant taxa and color preferences among the recorded lepidopteran visitors (Balfour et al., 2022).

## RESULTS

### Checklist of Lepidopteran Visitors and Flowering Plants

As of 27 November 2025, the “Lepidoptera in Metropolitan Manila” community project on iNaturalist comprised 1,339 research-grade observations representing 149 species. From this dataset, 126 research-grade records corresponding to 38 lepidopteran taxa (including species and subspecies) were selected for analysis in the present study (Table 1). Butterfly visitors were dominated by members of the families Pieridae (n = 38), Papilionidae (n = 30), and Nymphalidae (n = 31). Only three Philippine endemic species were recorded: *Pareronia boebera*, *Troides rhadamantus*, and *Ypthima stellera*. Four moth species were also documented (n = 4), namely *Amata heubneri*, *Nyctemera coleta* (Erebidae), *Spoladea recurvalis* (Crambidae), and *Ophthalmis lincea* (Noctuidae).

**TABLE 1.**
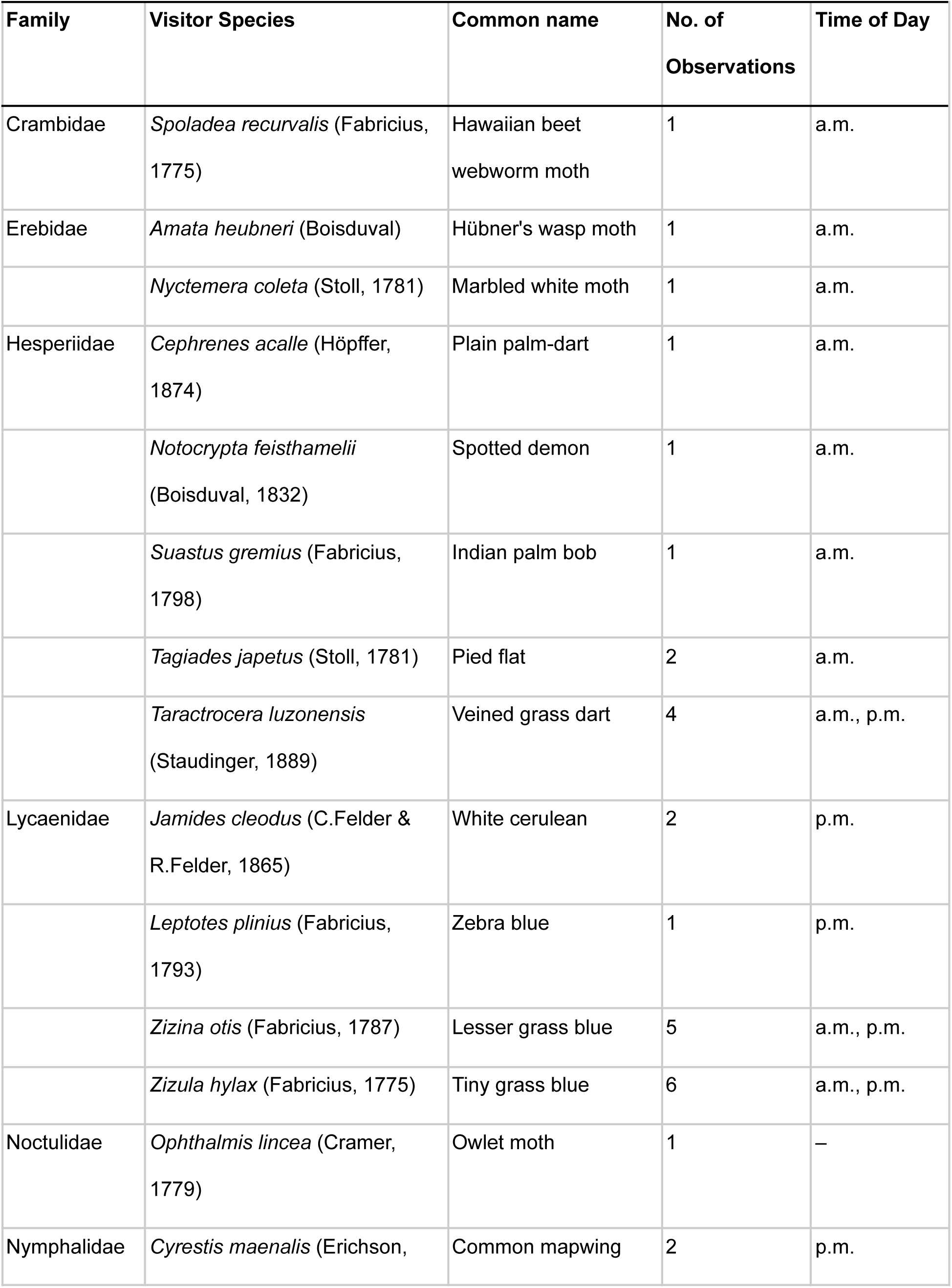
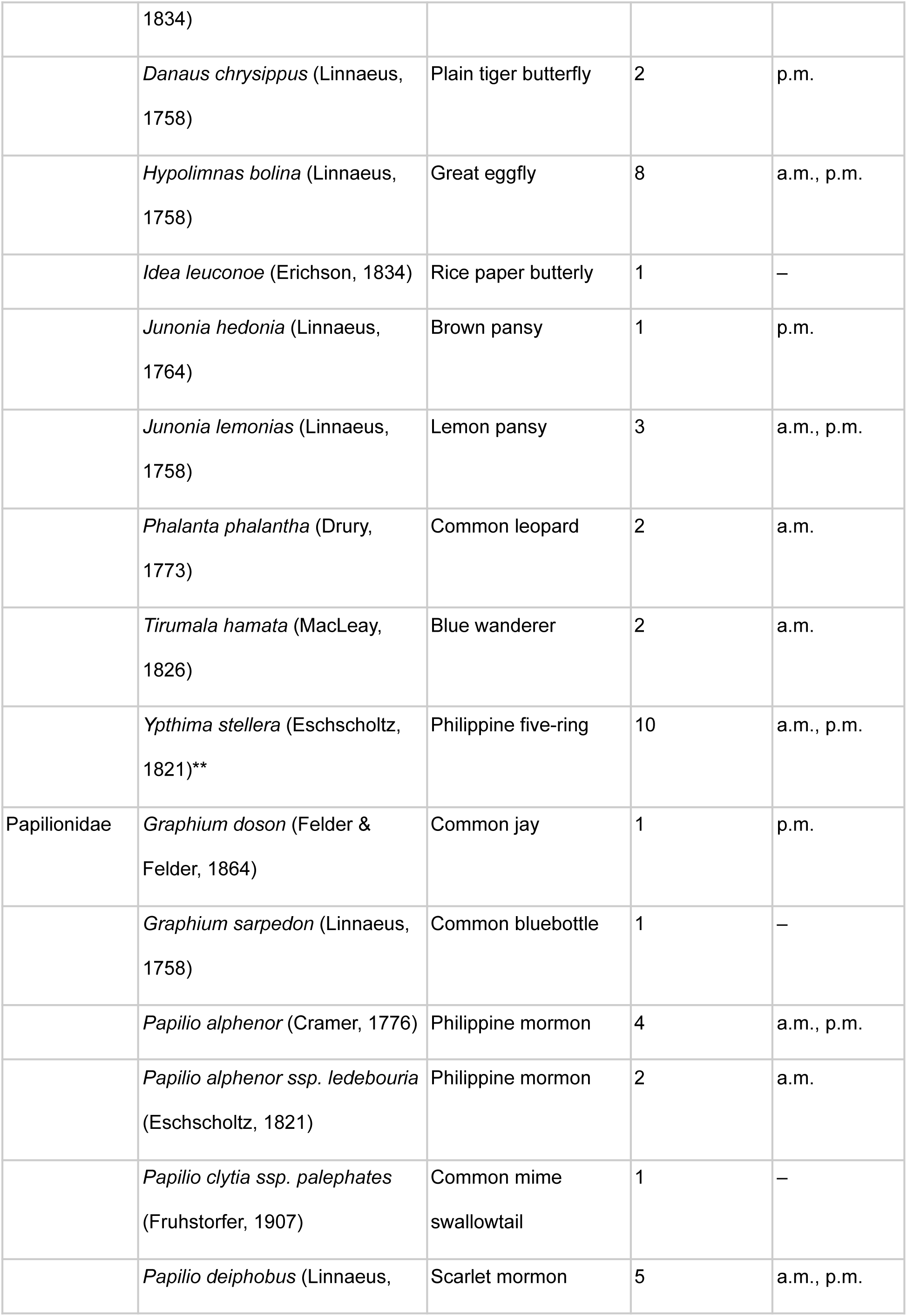
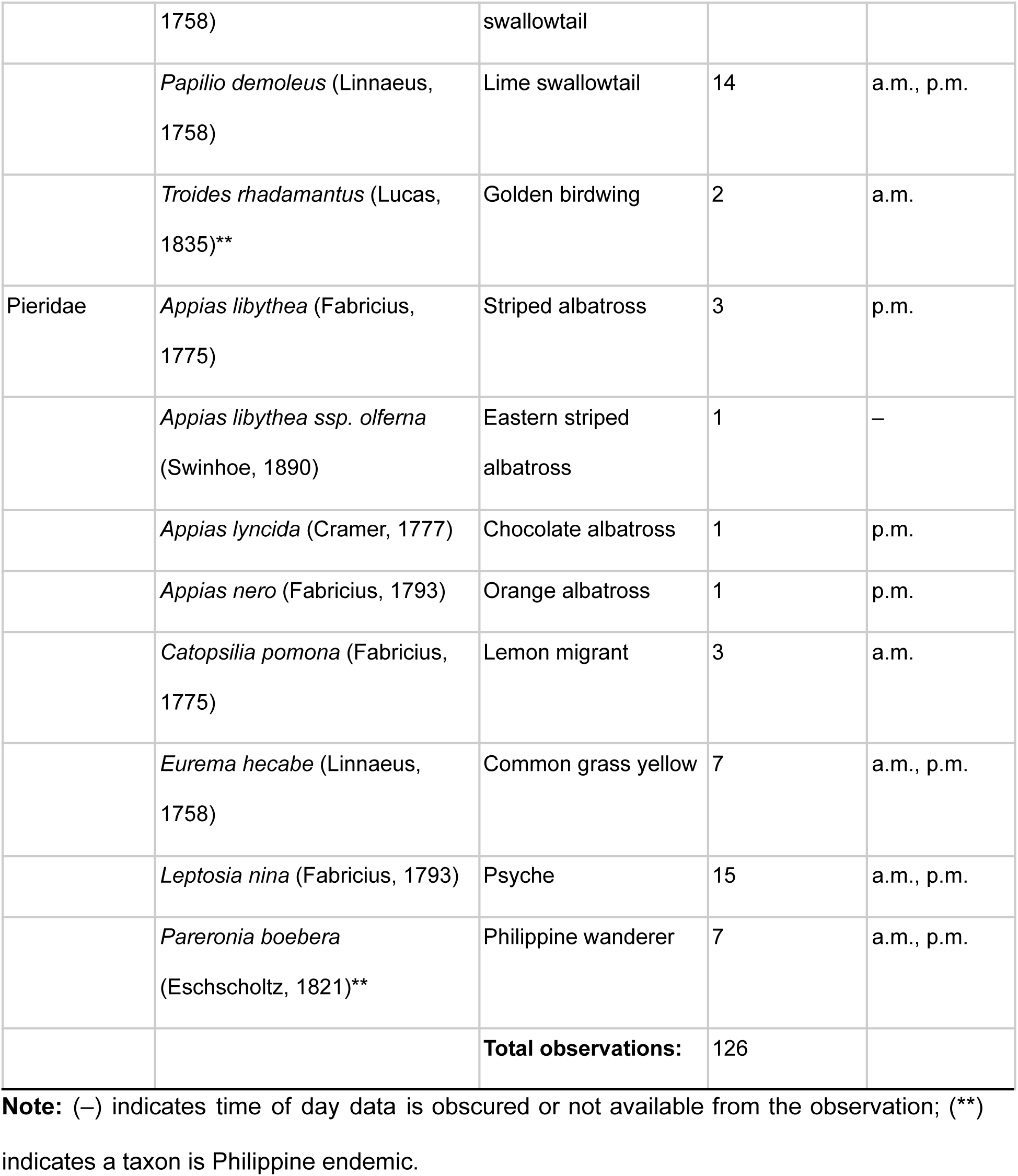
Lepidopteran visitors documented in Metro Manila by iNaturalist users from September 2015 to November 2025.

A total of 49 plant taxa visited by lepidopterans were identified to the genus, species, and cultivar levels (Table 2). The most frequently visited plant families were Asteraceae (n = 32), represented by common weedy species such as *Tridax procumbens* as well as cultivated ornamentals like *Cosmos* and *Sphagneticola*, and Rubiaceae (n = 20), which included the widely cultivated ornamental *Ixora* and the native ornamental *Mussaenda*. Visited plants were predominantly exotic (n = 97), with notable weedy exotics including *Tridax procumbens* (n = 22) and *Lantana camara* (n = 10), and common ornamental exotics such as *Ixora* spp. (n = 13), *Catharanthus roseus* (n = 5), and *Duranta erecta* (n = 5). Only 25 records involved native plant species, primarily *Alternanthera sessilis* (n = 9) and ornamental *Mussaenda* spp. (n = 3). Presented in Figures 4 and 5 are some images of representative Lepidoptera visiting flowers in Metro Manila.

**TABLE 2.**
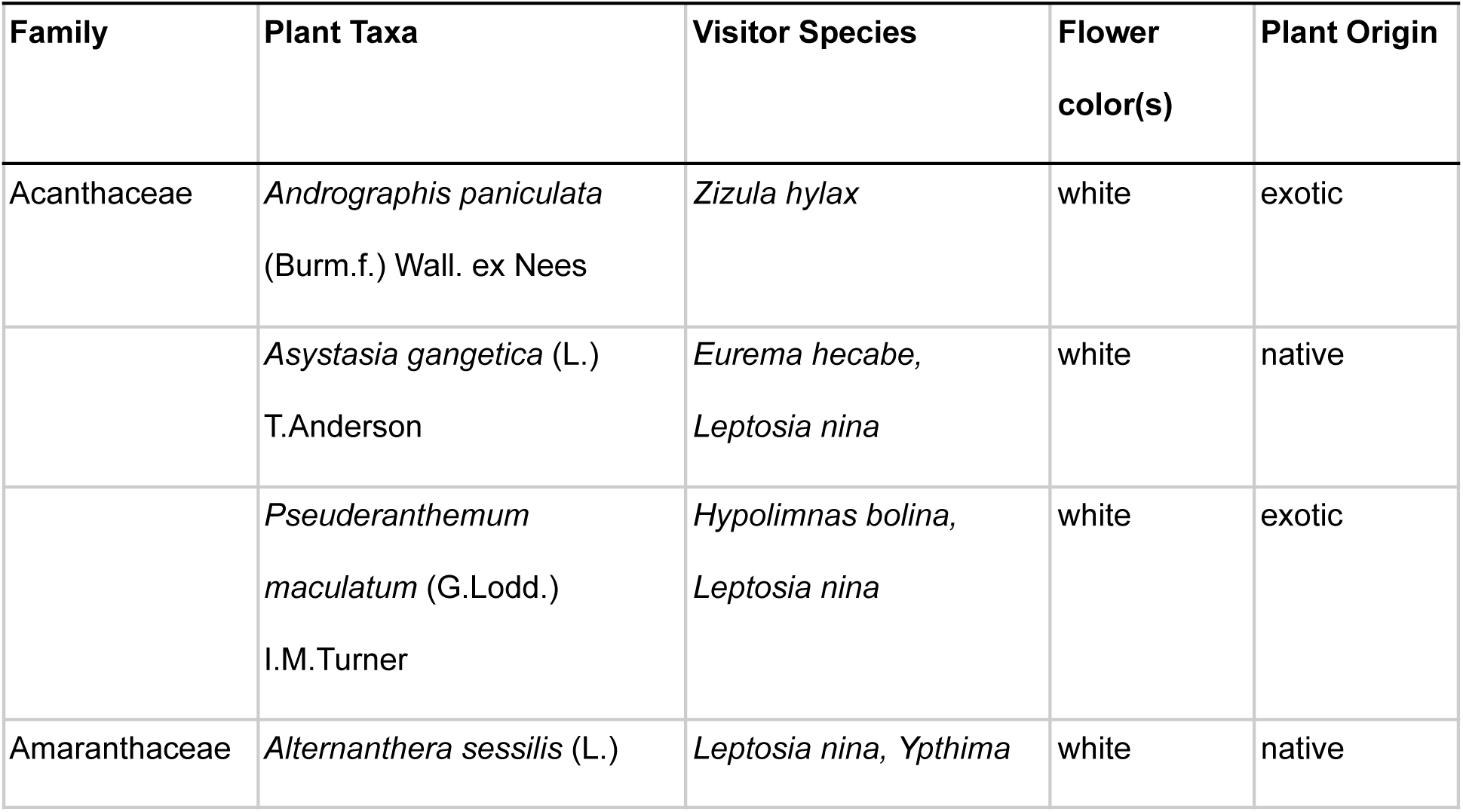
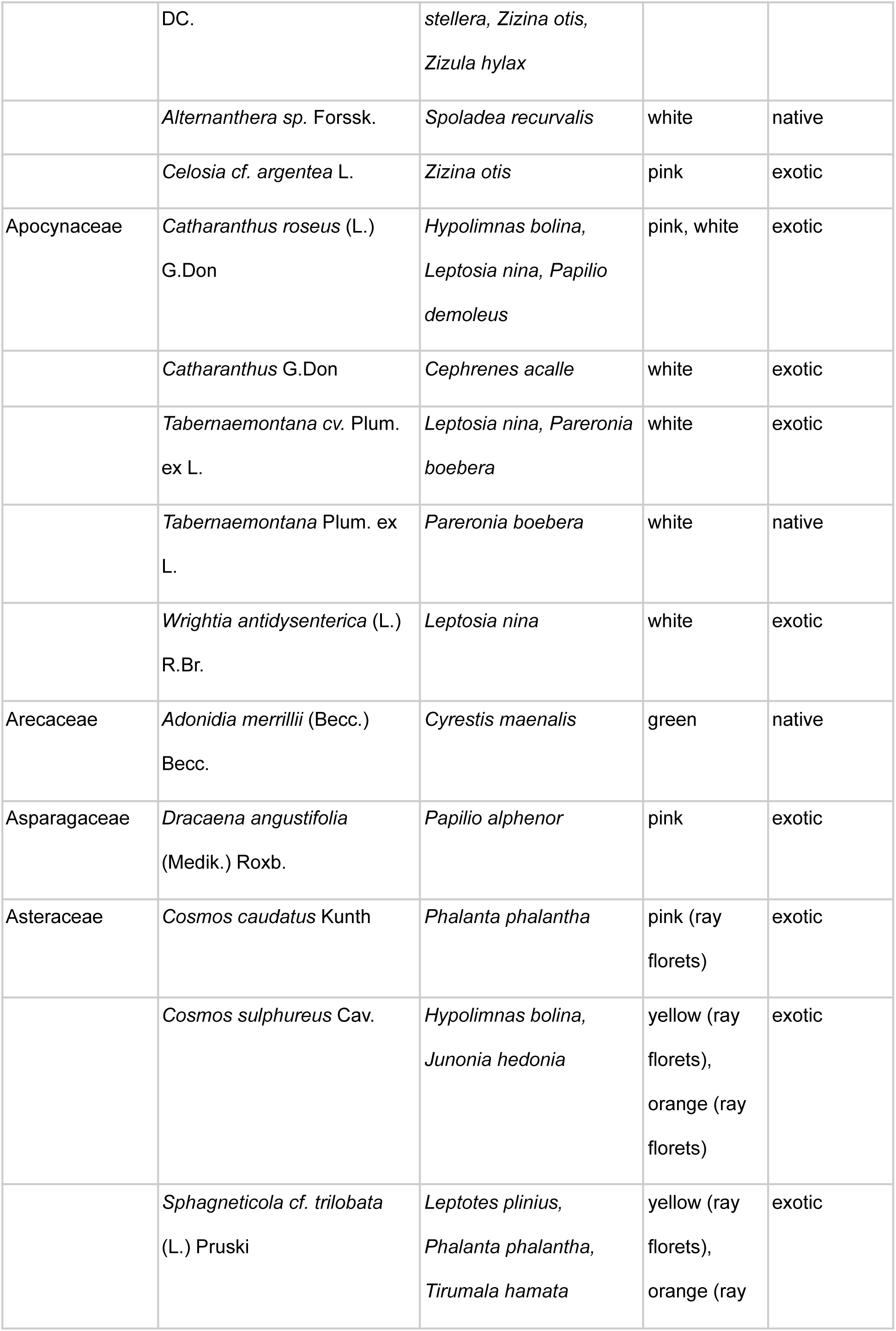
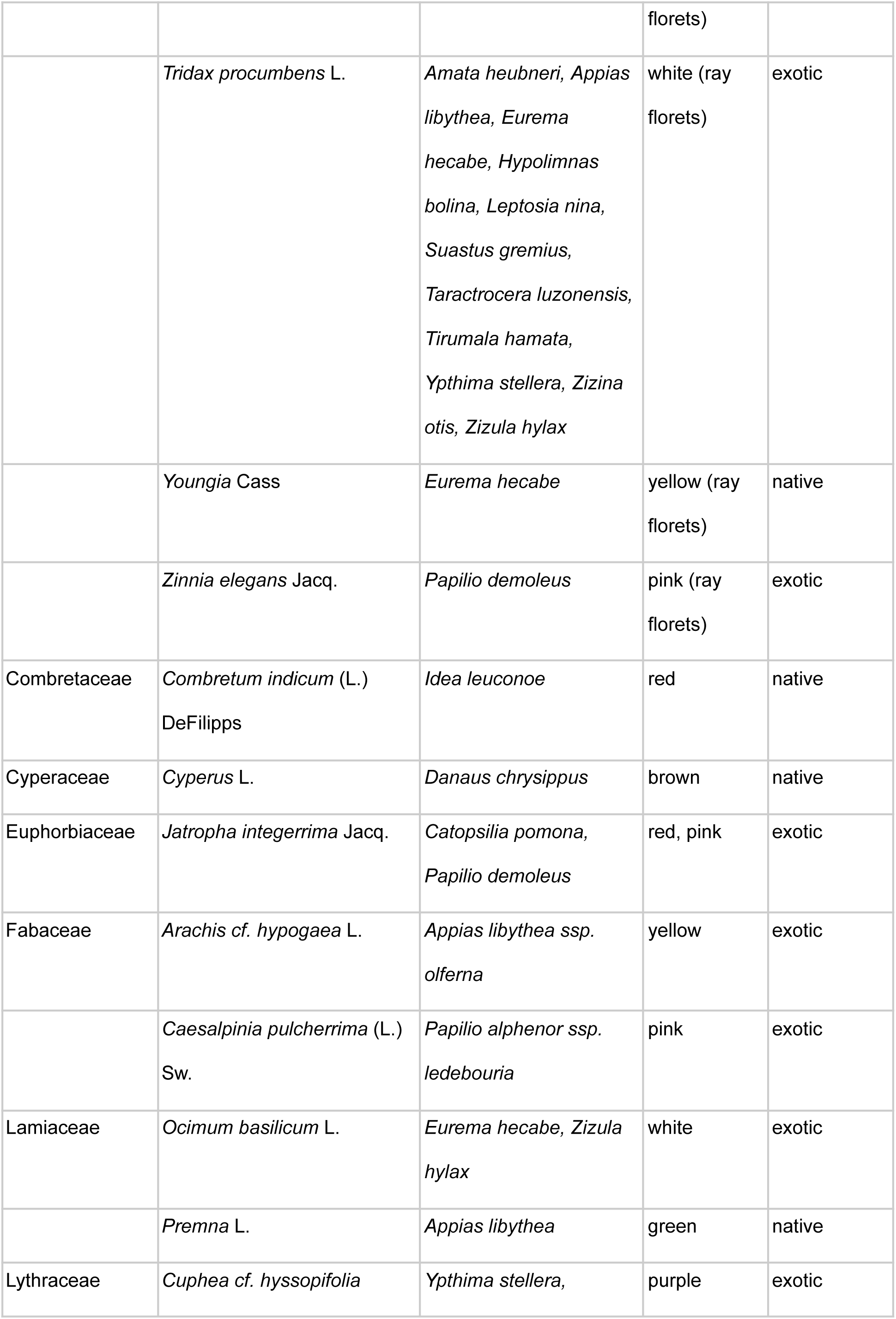
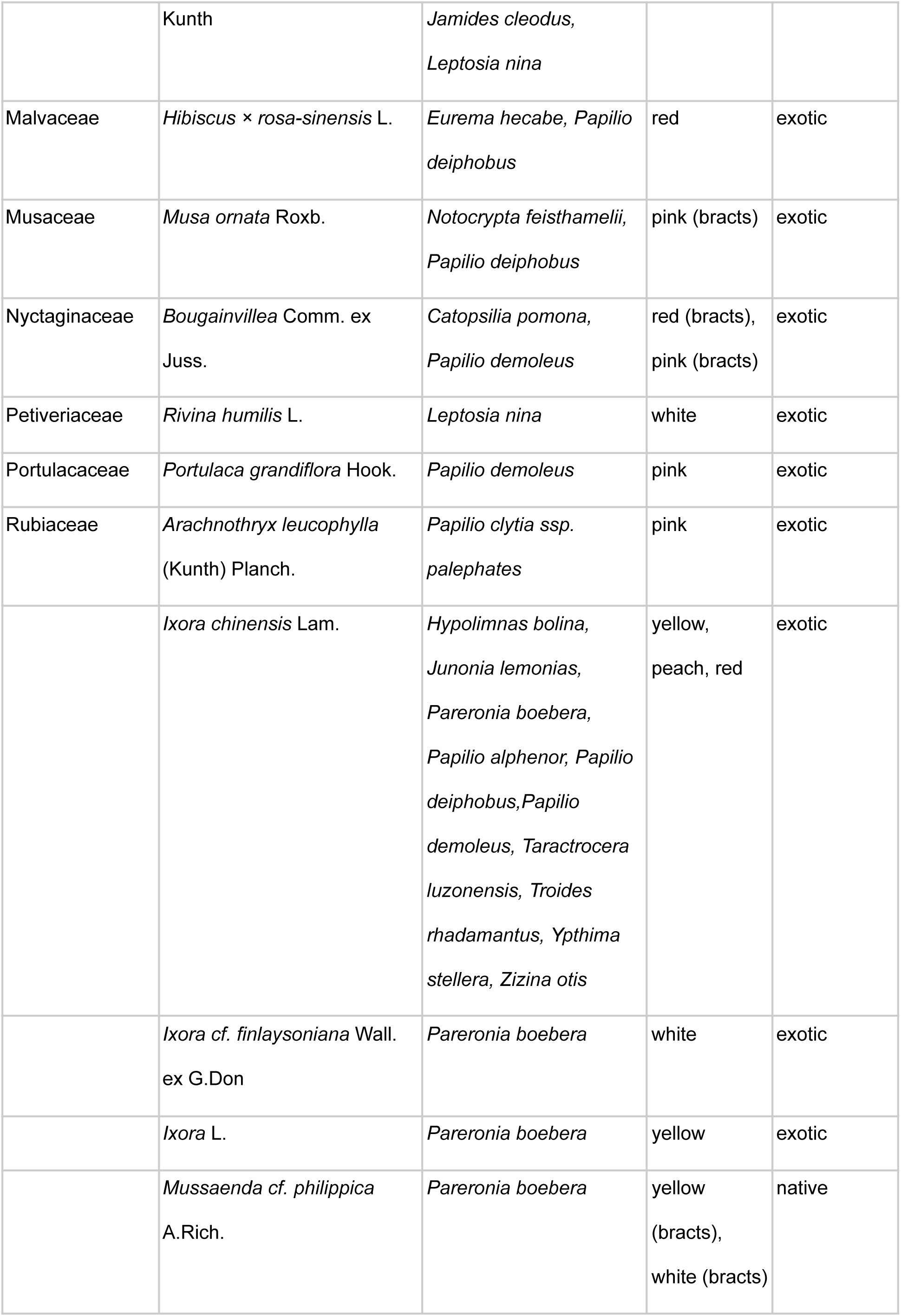
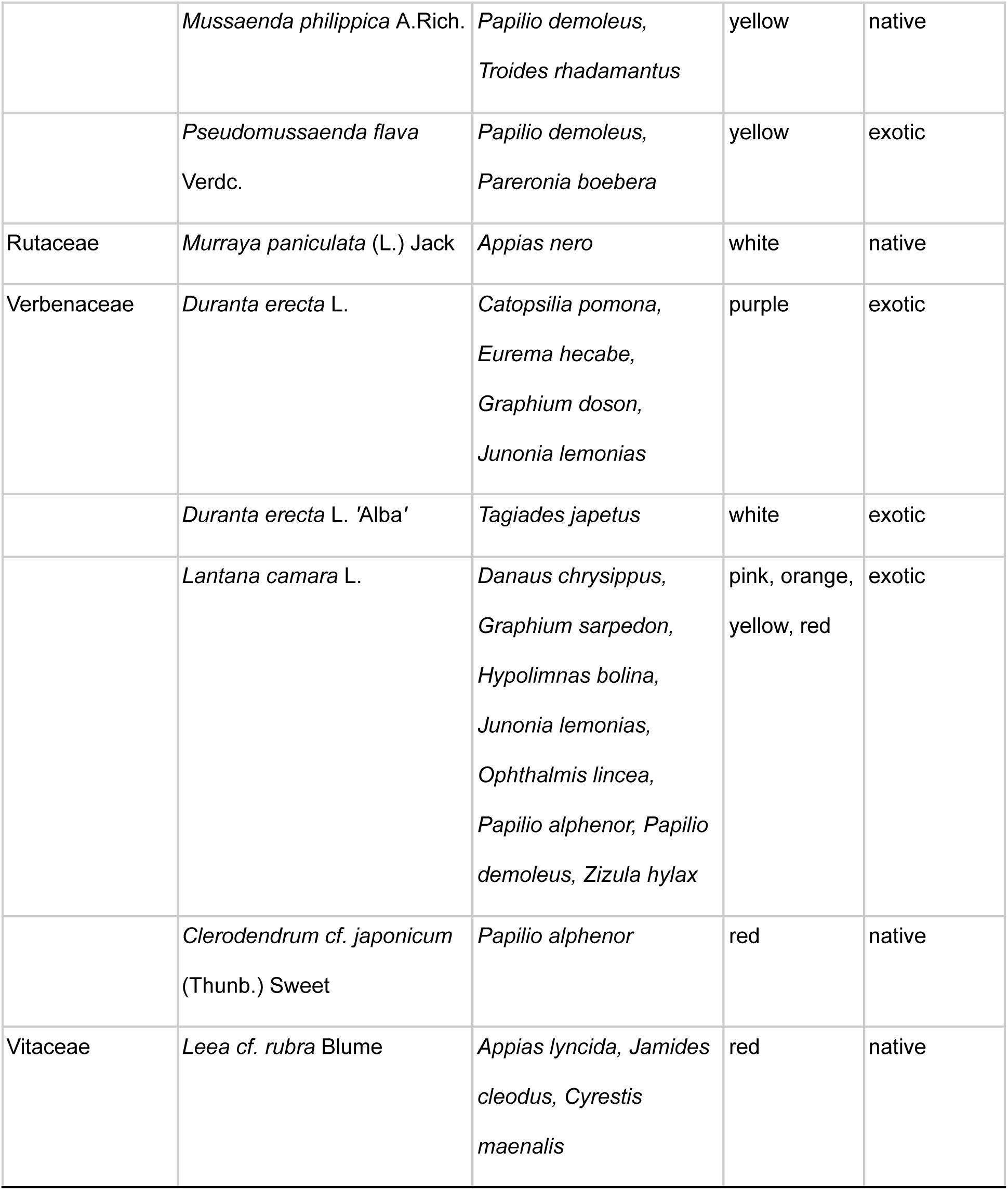
List of flowering plants visited by lepidopteran species documented in Metro Manila by iNaturalist users from September 2015 to November 2025.

**Figure 2.**
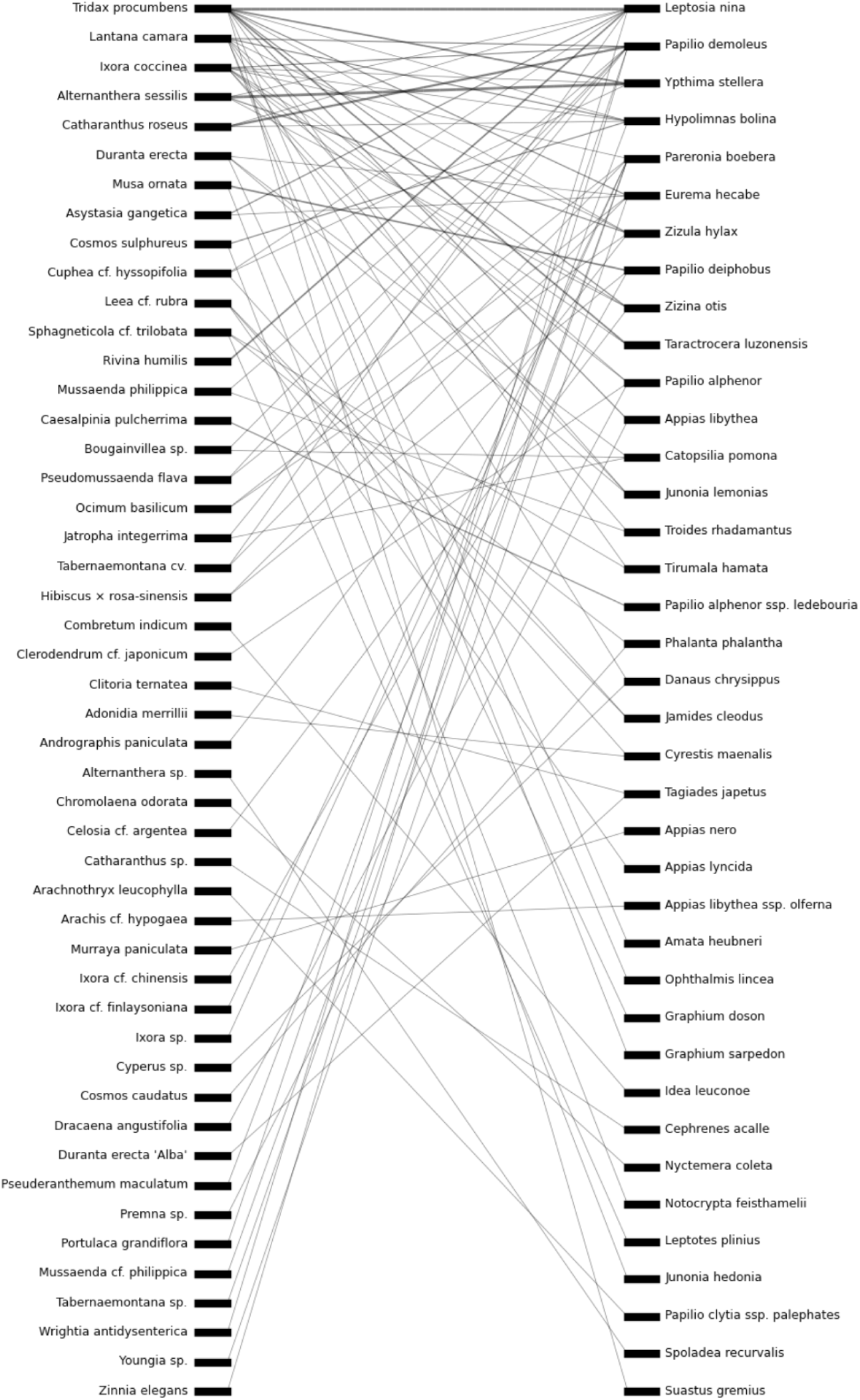
Bipartite network presenting Lepidopteran visitor-flowering species interaction in Metro Manila using citizen science data.

**Figure 3.**
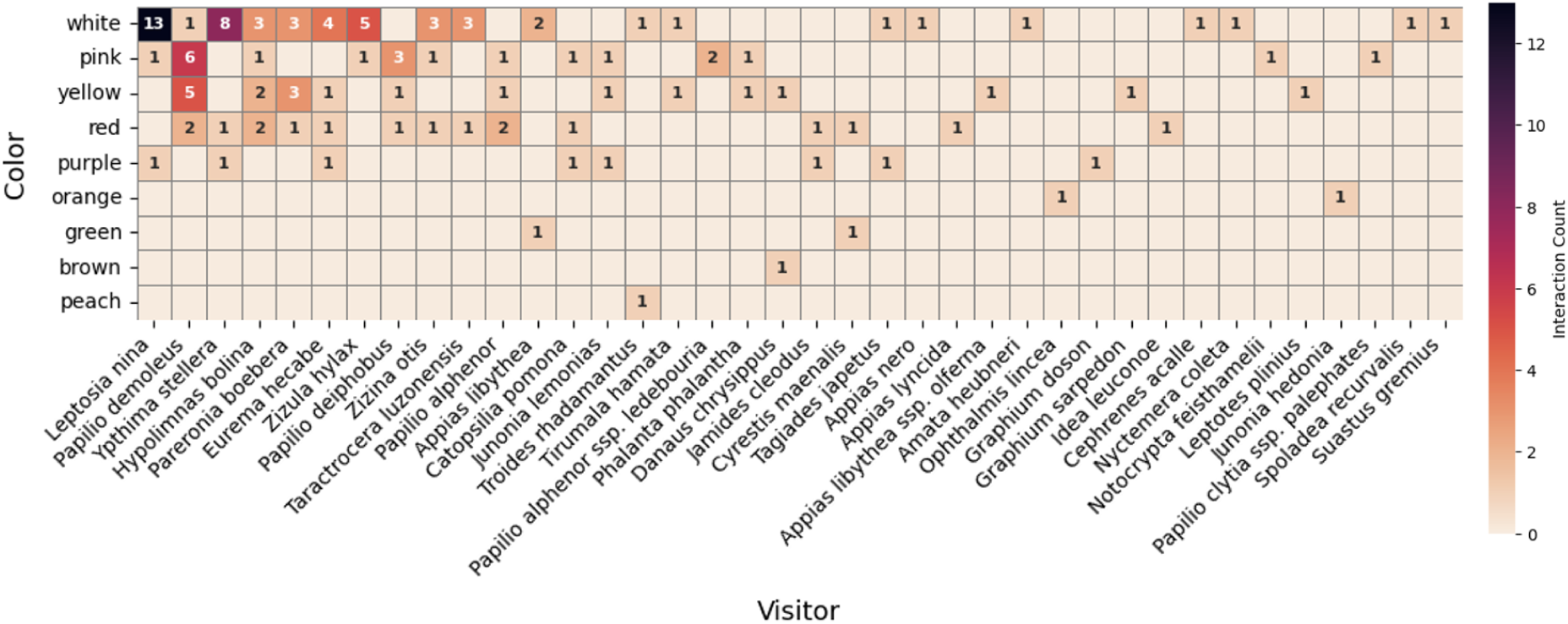
Flower Color-Lepidopteran Visitor Interaction Heatmap

**Figure 4.**
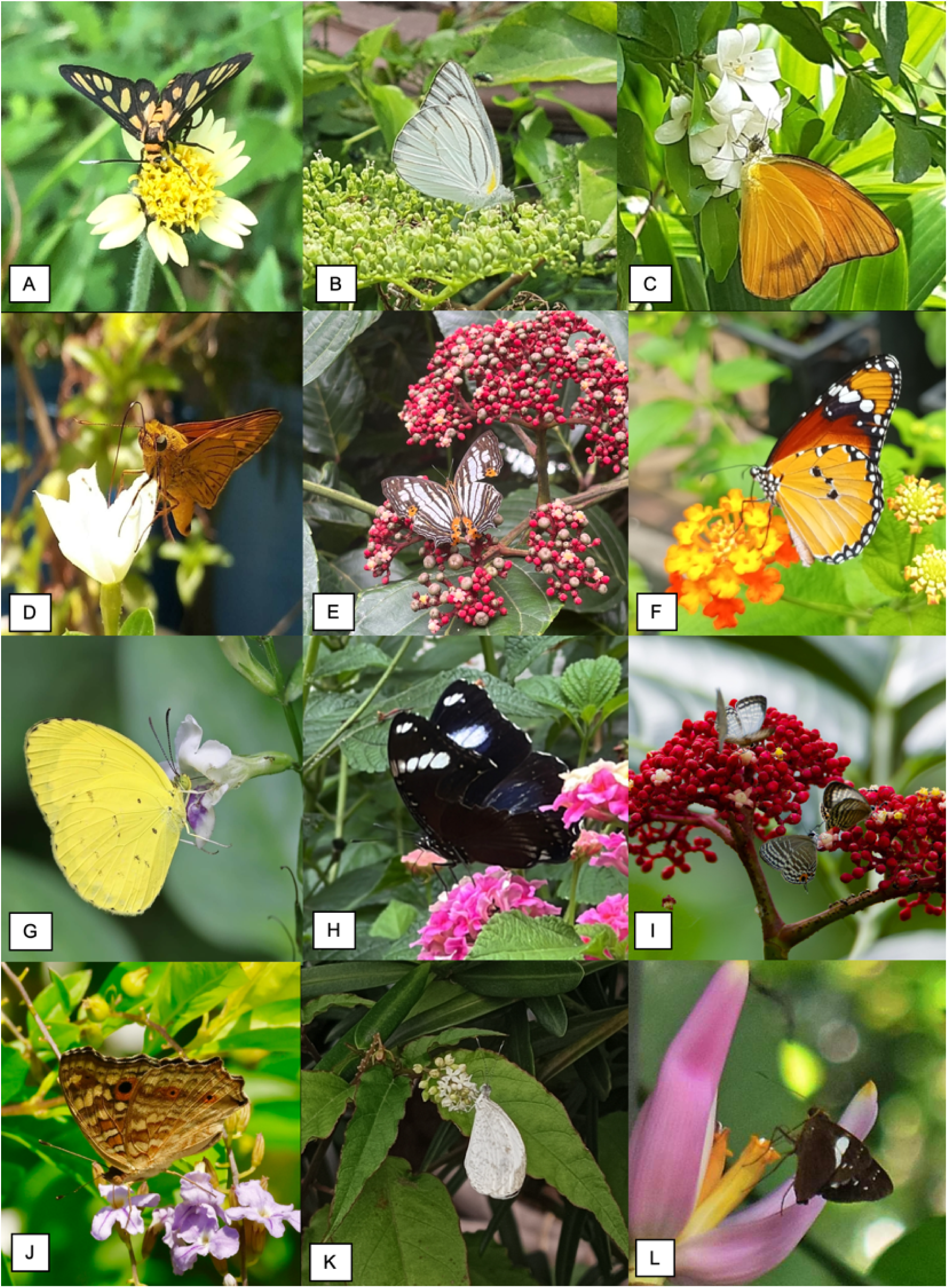
Representative Lepidoptera visiting flowers in Metro Manila showing interactions between selected species (Images from iNaturalist): (A) *Amata heubneri* and *Tridax procumbens* (© rossi107) (B) *Appias libythea* and *Premna sp.* (© mjbirder) (C) *Appias nero* and *Murraya paniculata* (© aniruddhachatterjee) (D) *Cephrenes acalle* and *Catharanthus roseus* (© wilhelmtan) (E) *Cyrestis maenalis* and *Leea cf. rubra* (© piper_andrea) (F) *Danaus chrysippus* and *Lantana camara* (© lnuacti) (G) *Eurema hecabe* and *Asystasia gangetica* (© mjbirder) (H) *Hypolimnas bolina* and *Lantana camara* (© joaquinduavit) (I) *Jamides cleodus* and *Leea cf. rubra* (© birdingphilippines) (J) *Junonia lemonias* and *Duranta erecta* (© birdexplorers) (K) *Leptosia nina* and *Rivina humilis* (© wilhelmtan) (L) *Notocrypta feisthamelii* and *Musa ornata* (© wilhelmtan).

**Figure 5.**
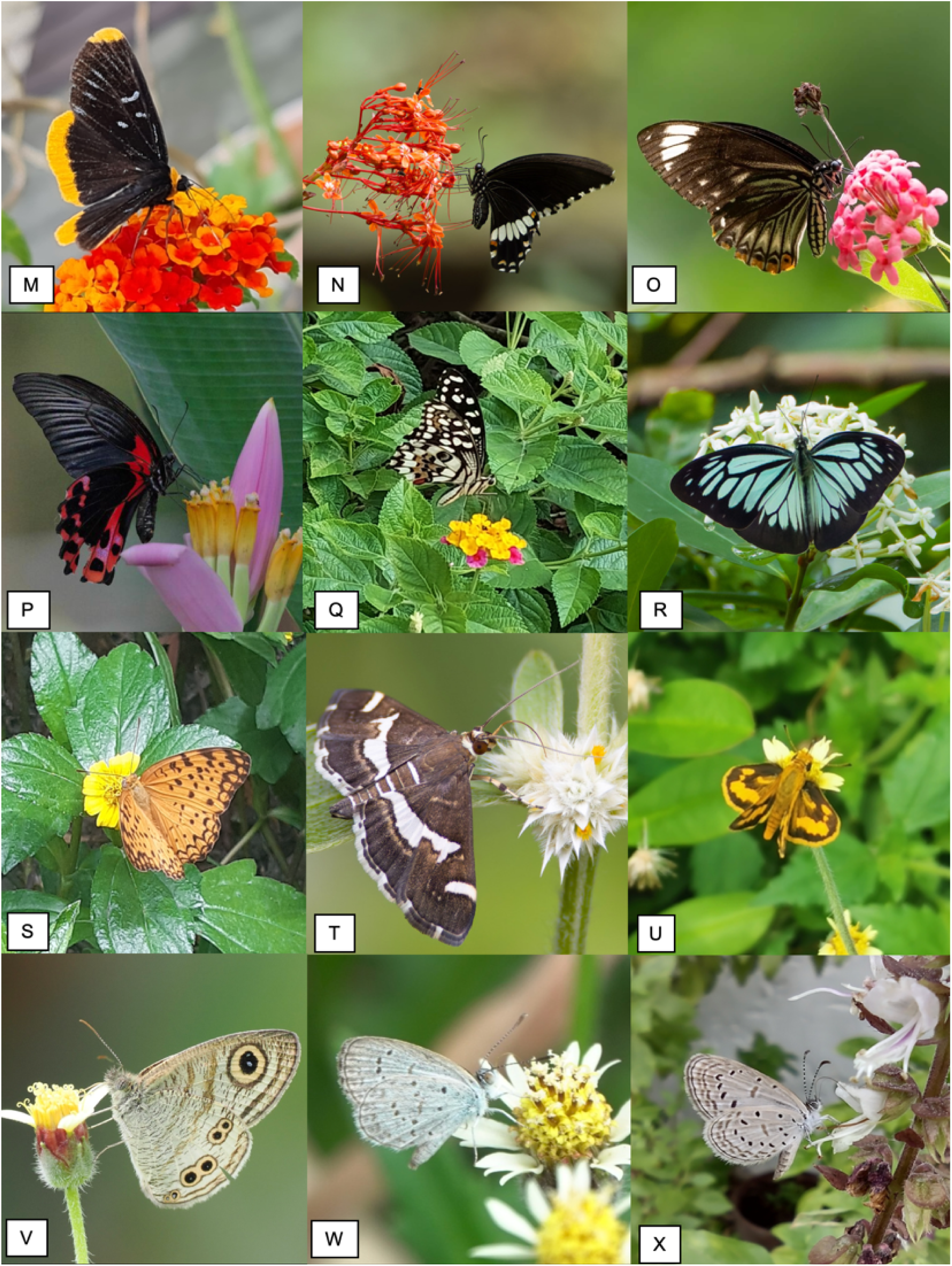
(Continued) (M) *Ophthalmis lincea* and *Lantana camara* (© popsie1) (N) *Papilio alphenor* and *Clerodendrum cf. japonicum* (© birdingphilippines) (O) *Papilio clytia ssp. palephates* and *Arachnothryx leucophylla* (© birdingphilippines) (P) *Papilio deiphobus* and *Musa ornata* (© ou_wildlife) (Q) *Papilio demoleus* and *Lantana camara* (© maylopez) (R) *Pareronia boebera* and *Ixora cf. finlaysoniana* (© birdingphilippines) (S) *Phalanta phalantha* and *Sphagneticola cf. trilobata* (© aniruddhachatterjee) (T) *Spoladea recurvalis* and *Alternanthera sp.* (© birdingphilippines) (U) *Taractrocera luzonensis* and *Tridax procumbens* (© yushe) (V) *Ypthima stellera* and *Tridax procumbens* (© mjbirder) (W) *Zizina otis* and *Tridax procumbens* (© kenny_well) (X) *Zizula hylax* and *Ocimum basilicum* (© bearbeer)

Spatially, most observations were recorded from Quezon City (n = 67), particularly within the Ateneo de Manila University campus, the University of the

Philippines–Diliman, and La Mesa Ecopark (Table 3). Temporally, the majority of records (n = 94) were collected during the wet season (June–November). The earliest observation included in this study dates to September 2015, while most records (n = 110) were submitted between 2020 and 2025.

**TABLE 3.**
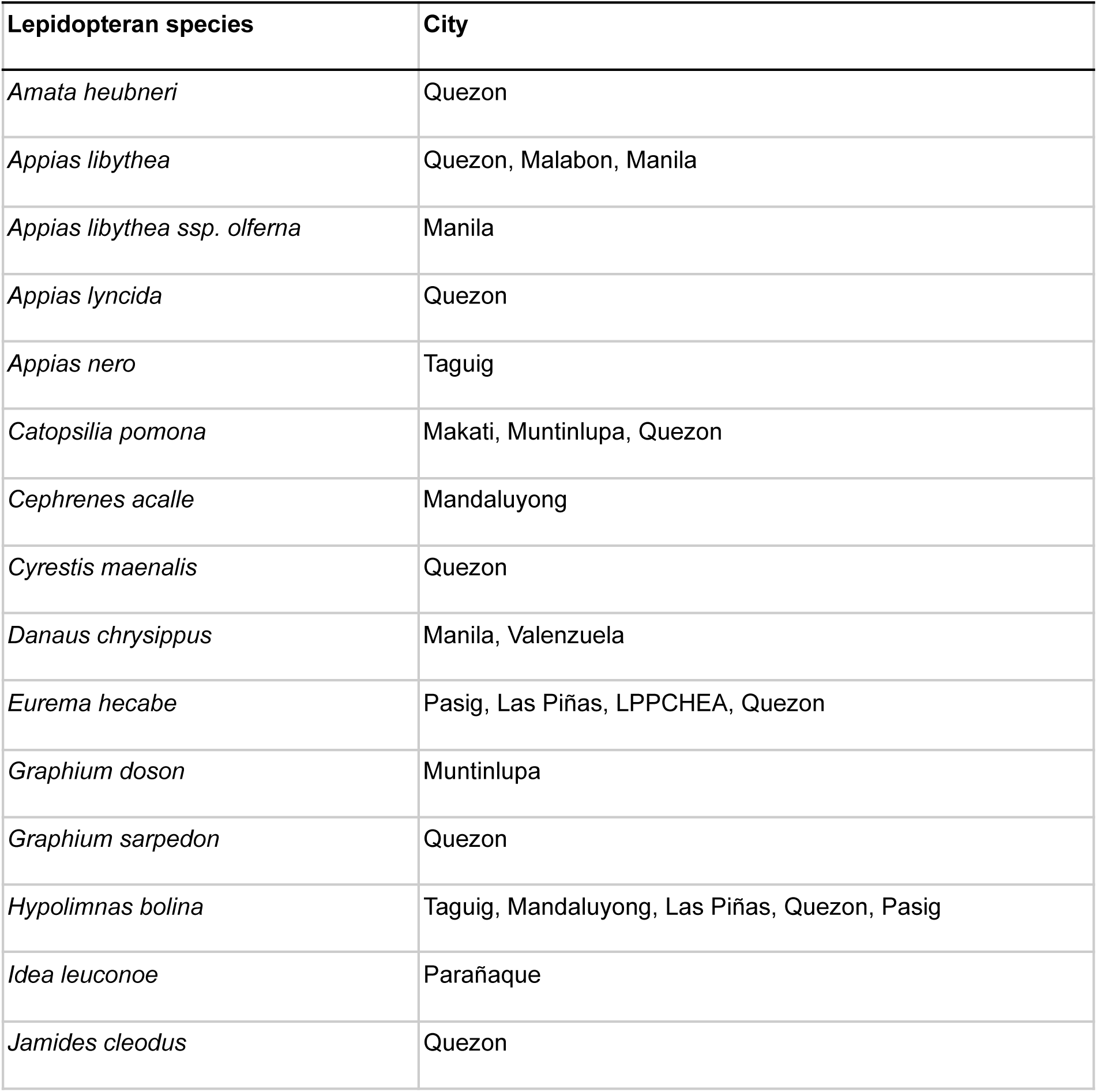
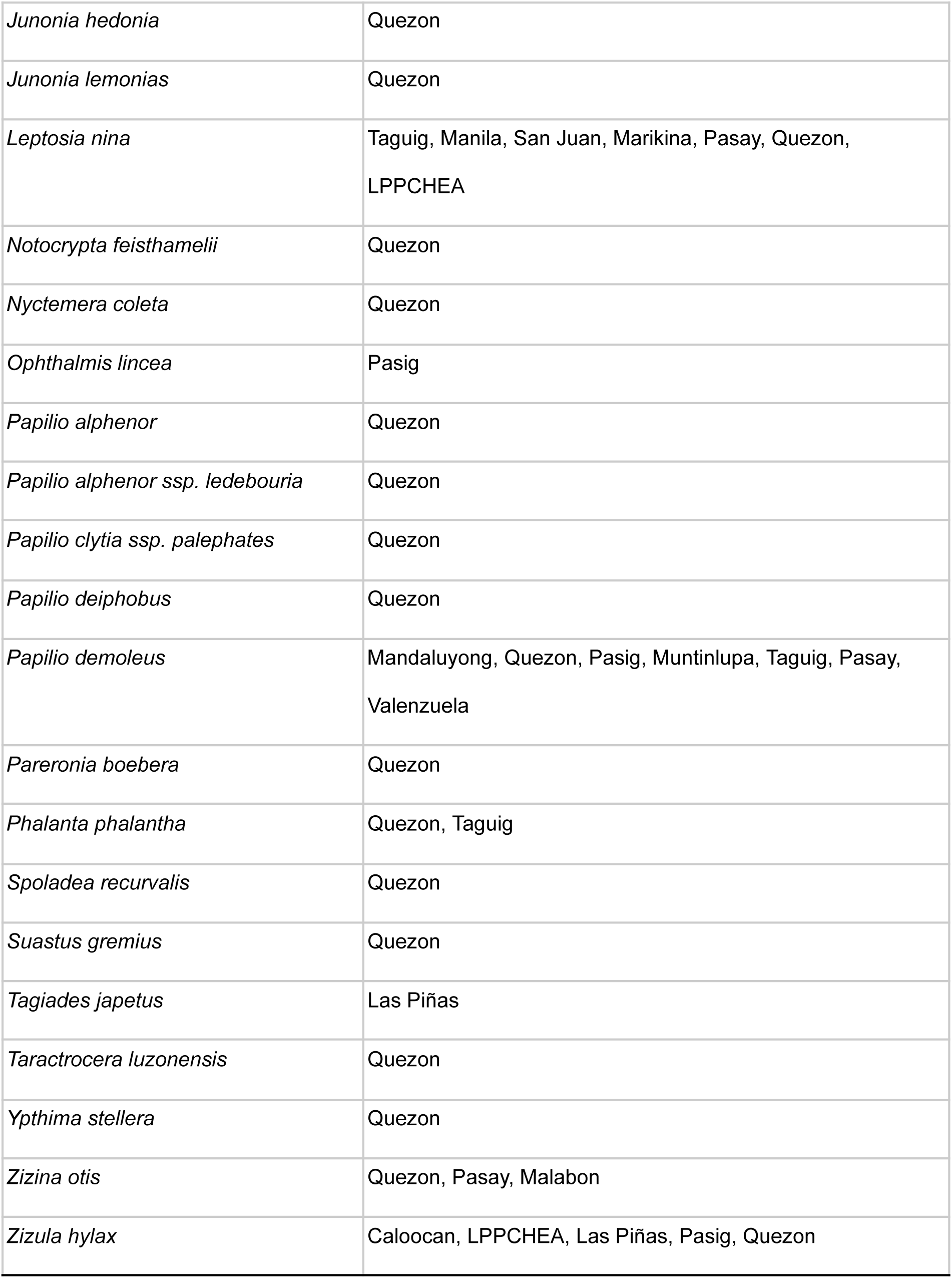
Lepidopteran species recorded in Metro Manila and the corresponding cities/areas where they were documented on iNaturalist from September 2015 to November 2025.

### Lepidopteran visitor-flowering species bipartite network

The resulting bipartite network (Fig. 2) illustrates all recorded flower–visitor interactions. The network exhibits a generalized interaction structure, with most butterflies visiting multiple flowering plant species and many plants being associated with more than one lepidopteran visitor. Several species occupied central positions within the network due to their high degree of interaction. Among the lepidopteran visitors, *Papilio demoleus* (Papilionidae), *Leptosia nina* (Pieridae)*, Ypthima stellera* (Nymphalidae), *Hypolimnas bolina* (Nymphalidae), and *Eurema hecabe* (Pieridae) showed the highest number of recorded plant associations. On the plant side, frequently visited species are *Tridax procumbens* (Asteraceae), *Lantana camara* (Verbenaceae), *Ixora coccinea* (Rubiaceae), *Alternanthera sessilis* (Amaranthaceae), and *Catharanthus roseus* (Apocynaceae). These species served as major hubs linking multiple lepidopteran taxa.

Most species on both trophic levels displayed relatively few interactions, reflected by numerous thin or isolated links in the network. Several butterflies, such as *Suastus gremius* (Hesperiidae), *Spoladea recurvalis* (Crambidae), and *Papilio clytia ssp. palephates* (Paplionidae), and several plant species such as *Zinnia elegans* (Asteraceae)*, Youngia sp.* (Asteraceae), and *Wrightia antidysenterica* (Apocynaceae) appeared only once or twice in the dataset, contributing to a long tail of low-frequency interactions. The distribution of link density indicates that interactions were unevenly represented, with a few common species dominating most of the network connections.

Overall, the bipartite visualization summarizes the full set of documented interactions between flowering plants and lepidopteran visitors within Metro Manila using citizen science data, providing a clear depiction of both frequently occurring and infrequent associations.

### Lepidopteran visitors and preferences on flower colors

The flower color–lepidopteran visitor interaction heatmap (Figure 3) shows a clear dominance of interactions on white, yellow, and pink flowers across the recorded lepidopteran taxa. White flowers exhibited the highest overall interaction frequencies, with particularly strong visitation by *Leptosia nina*, *Papilio demoleus*, *Ypthima stellera*, and *Hypolimnas bolina*. Yellow flowers also received substantial visitation, especially from pierid butterflies such as *Eurema hecabe*, *Catopsilia pomona*, *Appias lyncida*, and *Appias libythea*. Pink flowers showed moderate interaction levels, involving primarily *Pareronia boebera*, *Hypolimnas bolina*, and *Papilio polytes*.

Red and purple flowers registered comparatively fewer interactions and were visited by a limited subset of taxa, while orange, green, brown, and peach flowers recorded only isolated visitation events. Moth species, including *Nyctemera coleta*, *Spoladea recurvalis*, and *Ophthalmis lincea*, were predominantly associated with white and pale-colored flowers. High interaction values were concentrated among widespread generalist butterflies such as *Leptosia nina*, *Papilio demoleus*, and *Appias* spp., whereas several taxa exhibited low, sporadic visitation across multiple flower colors.

## DISCUSSION

### Visitor-flower interactions

The visitor-flower interactions documented in this study reveal a predominantly generalized lepidopteran assemblage in Metro Manila. The bipartite network is characterized by several highly connected species on both the plant and butterfly sides, consistent with patterns often observed in urbanized landscapes where floral resources are patchy, dominated by ornamentals, and continuously replenished across seasons. Commonly cultivated or weedy, naturalized species such as *L. camara*, *T. procumbens*, *I. coccinea*, and *C. roseus* formed the principal floral hubs supporting multiple butterfly taxa. These plants are widely distributed in gardens, roadsides, and small green spaces, allowing them to act as persistent nectar sources throughout the year (Mukherjee et al., 2015; Sze Huei, 2024).

Similarly, the most frequently observed butterflies—*Papilio demoleus*, *Hypolimnas bolina*, *Eurema hecabe*, and *Junonia lemonias*—are disturbance-tolerant, highly mobile, and capable of exploiting diverse nectar sources. Their dominance in the network reflects their ubiquity in the urban matrix and their capacity to utilize both native and introduced plants. The diffuse structure of the network, with numerous low-frequency interactions and few specialists, aligns with the expected composition of heavily modified tropical cities, where habitat fragmentation and continuous human disturbance favor ecological generalists (Muñoz-Galicia et al., 2023; Ancilloto et al., 2024).

Despite its generality, the network also highlights the presence of less frequently encountered taxa and rare interaction pairs. Species appearing only once or twice in the dataset, whether due to genuine rarity or limited documentation, demonstrate the potential of citizen-science platforms to detect interactions that may be overlooked in systematic surveys. Such incidental records provide valuable preliminary insights into the nectar resources of endemic or native butterflies and the potential role of native or less conspicuous plants in sustaining urban insect diversity (Braman & Griffin, 2022).

This study documents new or noteworthy nectar associations for three Philippine endemic butterflies—*Pareronia boebera*, *Troides rhadamantus*, and *Ypthima stellera*—based on opportunistic citizen-science records from Metro Manila. These observations provide valuable natural history information for species that are seldom recorded feeding in urban environments and highlight the role of cultivated and spontaneous plants in sustaining endemic Lepidoptera within the metropolitan landscape.

The Philippine wanderer (*Pareronia boebera*) was recorded visiting multiple ornamental species within the Rubiaceae and Apocynaceae. Confirmed floral associations include *Ixora cf. finlaysoniana*, *Ixora coccinea*, *Ixora* sp., *Mussaenda cf. philippica*, *Pseudomussaenda flava*, *Tabernaemontana* cv., and *Tabernaemontana* sp. Several of these—particularly *Pseudomussaenda flava*, *Mussaenda cf. philippica*, *Ixora cf. finlaysoniana*, and the *Tabernaemontana* records—do not appear in existing literature and therefore represent new or previously unreported nectar associations for this endemic species. Mape et al. (2022) documented four nectar plants for *P. boebera*: *I. coccinea*, *Plumbago auriculata*, *Turnera ulmifolia*, and *Bougainvillea spectabilis*. Of these, only *I. coccinea* overlaps with the present findings. All other plant associations reported in this study—*Ixora cf. finlaysoniana*, *Ixora* sp., *Mussaenda cf. philippica*, *Pseudomussaenda flava*, *Tabernaemontana* cv., and *Tabernaemontana* sp.—constitute additional nectar records to Mape et al.’s (2022) report.

The golden birdwing (*Troides rhadamantus*), a charismatic Philippine endemic, was documented nectaring on *Ixora coccinea* and *Mussaenda philippica*. Although birdwings are broadly known to visit ornamental flowers, specific records from Metro Manila remain limited. These observations underscore the role of nectar-rich ornamental shrubs—particularly *Ixora* and *Mussaenda*—in sustaining adult foraging activity of this large, wide-ranging species within urban and suburban landscapes. Mape et al. (2022) reported several nectar plants associated with *T. rhadamantus*, including *Murraya paniculata*, *Premna odorata*, *Lantana camara*, *Duranta erecta*, *Catharanthus roseus*, *Wrightia antidysenterica*, *Kopsia cf. fruticosa*, and *Adenium obesum*. The present study adds *Mussaenda philippica* as an additional nectar record for *T. rhadamantus*, expanding the documented range of floral resources used by this species in metropolitan habitats.

The Philippine five-ring (*Ypthima stellera*) showed repeated visitation to *Alternanthera sessilis*, with multiple independent observations providing strong evidence that this widespread ruderal herb serves as a regular nectar resource in disturbed or roadside habitats. Additional interactions were documented with *Cuphea cf. hyssopifolia*, *Ixora coccinea*, and *Tridax procumbens*, indicating a broader range of nectar sources than previously reported for the species. Although *Y. stellera* is common in grasslands and open urban spaces, its nectar associations remain poorly characterized in literature. Mape et al. (2022) listed six nectar plants for the species—*Premna odorata*, *T. procumbens*, *Bidens pilosa*, *Turnera ulmifolia*, *Jatropha podagrica*, and *Russelia equisetiformis*. The present study contributes additional records, specifically its use of *Cuphea cf. hyssopifolia*, *Ixora coccinea*, and *A. sessilis*, most of which are abundant in residential gardens, parks, and roadside vegetation in Metro Manila. These new nectar associations suggest that *Y. stellera* readily exploits both cultivated ornamentals and naturalized weeds, further supporting its characterization as a flexible, disturbance-tolerant species. Its repeated use of *A. sessilis* highlights the ecological importance of ruderal flora in sustaining butterfly populations within highly modified landscapes. Likewise, interactions with *Cuphea* and *Ixora* expand its known nectar repertoire, while the records of *T. procumbens* are consistent with the generalist feeding patterns commonly observed among nymphalids in urban mosaics.

The use of opportunistic citizen-science observations, however, introduces inherent limitations. Records are influenced by observer effort, accessibility of locations, photogenic species, and uneven sampling across seasons and barangays (Johnston et al., 2023; Pernat et al., 2024). Consequently, link frequency does not necessarily equate to true interaction strength or ecological importance (Dickinson et al., 2010). Plant species located in private gardens, gated communities, or restricted areas may be underrepresented. Conversely, widespread ornamentals may appear disproportionately important due to their visibility and accessibility. These biases underscore that while opportunistic data can reveal the breadth of possible interactions, it cannot fully quantify interaction rates or specialization without complementary systematic field sampling (Jacquemin et al., 2020).

Nevertheless, the dataset demonstrates the value of citizen science as a tool for documenting pollinator–plant interactions in highly urbanized contexts. The breadth of interactions captured—spanning both common and rarely photographed species—illustrates how digital platforms can help fill spatial and temporal gaps in biodiversity information, especially in cities where formal surveys are limited (Bitonto et al., 2025). The network generated here provides a baseline inventory of flower-visiting butterflies in Metro Manila and highlights plant species that currently sustain a wide range of Lepidoptera within fragmented urban green spaces.

Overall, this study contributes to the growing evidence that urban environments, despite intense anthropogenic pressure, can support a diverse set of butterfly species provided that floral resources remain available. The interaction patterns observed here emphasize the importance of maintaining nectar-rich gardens, roadside vegetation, and pockets of native flora as essential components of urban biodiversity conservation. Continued integration of citizen-science data with targeted ecological surveys will further enhance our understanding of pollinator dynamics and help guide habitat enhancement strategies within Metro Manila and other tropical cities.

### Insights into color preference of lepidopterans

The strong concentration of lepidopteran visitation on white and yellow flowers observed in this study is consistent with visual foraging preferences of diurnal butterflies for bright, high-reflectance floral cues that enhance detectability in complex urban environments (Dylewski, Mackowiak, & Banaszak-Cibicka, 2020; Ramirez-Restrepo & MacGregor-Fors, 2017). This pattern is especially pronounced among Pieridae, a family known for innate attraction to yellow flowers, which likely explains the high interaction frequencies recorded for *Eurema*, *Catopsilia*, and *Appias* species (Sinha et al., 2023). The overall dominance of white flowers further highlights their importance as primary nectar resources in Metropolitan Manila’s urban matrix.

The moderate but taxon-specific use of pink and red flowers suggests that these colors remain functionally important for certain butterflies such as *Pareronia boebera*, *Hypolimnas bolina*, and *Papilio polytes*. In contrast, the very low interaction frequencies observed for purple, orange, green, brown, and peach flowers likely reflect a combination of limited local availability of these floral colors in urban green spaces and observer sampling bias, rather than true ecological avoidance.

The tendency of moth species to concentrate on white and other pale-colored flowers is consistent with their crepuscular to nocturnal activity patterns, wherein high-contrast floral displays enhance visual detection under low-light conditions (Yurtsever, Okyar, & Guler, 2010; van der Kooi, 2021). The dominance of interactions by widespread generalist butterflies such as *Leptosia nina* and *Papilio demoleus* further reflects the characteristic structure of urban lepidopteran communities, which are typically shaped by habitat-tolerant species capable of exploiting abundant ornamental and weedy nectar sources.

The color–visitor associations observed in Metro Manila are also consistent with patterns reported in controlled behavioral studies. Sinha et al. (2023) demonstrated a strong butterfly preference for yellow flowers, with orange as a secondary option and lower visitation on white, red, pink, and violet. This pattern is partially mirrored in the present dataset, where butterflies most frequently interacted with white and yellow flowers, followed by orange and pink, whereas red and mixed-colored blooms received few observations. The prominence of white in our results likely reflects the high abundance of white-flowered weeds (e.g., Asteraceae) in urban habitats and their overrepresentation in citizen-science platforms rather than true behavioral preference.

Importantly, the presence of Philippine endemic butterflies (*Pareronia boebera*, *Troides rhadamantus*, and *Ypthima stellera*) among visitors to predominantly white, yellow, and pink flowers emphasizes the conservation value of maintaining nectar-rich, visually conspicuous flowering plants in highly urbanized landscapes. These floral colors are commonly associated with cultivated ornamentals and ruderal species such as *Tridax procumbens*, *Cosmos*, and *Ixora*, underscoring the role of urban planting practices in shaping pollinator visitation networks and potentially supporting both widespread and restricted-range species.

The use of citizen science data from iNaturalist provides a powerful platform for documenting large-scale patterns of plant–pollinator interactions across space and time, particularly in understudied megacities such as Metropolitan Manila. Its strengths lie in the high volume of verifiable, geo-referenced observations, broad taxonomic coverage, and the ability to capture rare or stochastic interaction events that are difficult to document through traditional field surveys alone. However, several limitations must also be recognized. Citizen science records are inherently subject to observer bias toward conspicuous species, accessible locations, and daylight activity periods, which may inflate the apparent importance of common butterflies and visually prominent flowers while underrepresenting cryptic taxa, nocturnal interactions, and less showy floral resources. Additionally, floral color assignments are based on photographic perception, which may vary with lighting conditions, camera settings, and observer interpretation.

Despite these limitations, the consistency of the dominant color-visitation patterns observed here with established pollination biology strongly supports the ecological validity of the results. When interpreted cautiously, citizen science–derived interaction data thus represent a valuable and cost-effective complement to traditional pollination studies, particularly for assessing broad-scale urban ecological patterns, guiding biodiversity-friendly urban greening strategies, and identifying candidate plant species for pollinator conservation in tropical megacities.

## CONCLUSION

While this study, leveraging opportunistic citizen-science data, successfully established the first flower-visitor network for Metro Manila—revealing a generalized butterfly community sustained by common urban ornamentals and ruderal plants—the findings must be interpreted with caution. Although new nectar associations were found for Philippine endemic species, the observed strong preference for white and yellow flowers likely reflects an amplification due to sampling bias, where citizen scientists naturally document interactions with the most abundant and easily visible floral resources. Therefore, to sustain urban butterfly diversity and inform true pollinator-friendly strategies, future efforts must focus on maintaining nectar-rich ornamentals, weedy plants, and native flora while strategically integrating targeted field surveys to validate and correct the biases of opportunistic monitoring.

## ACKNOWLEDGMENT

I extend my heartfelt gratitude to the iNaturalist community, whose publicly shared observations made this study possible. In particular, I acknowledge the invaluable contributions of the following users: acey_blue, aidan_franco, agdpineda, ajasuzano, anders_forsberg, aniruddhachatterjee, ashley_jade, bearbeer, bernardomichelle, billiedepnag, birdingphilippines, birdexplorers, carizacruz, charie, christinebbaldo, clarenceramos, clynysbllee, dngapala, ericbuado, henryedelman, jayvee1, jhayra_liam, jj24, jjstaana, joan_dumo, joaquinduavit, josh_cruz, joshuaguinto, juanmiguelsantos35, kenny_well, kenjiwota, komeilet, kwagirl_01, lazza, lnuacti, maylopez, meowa, mdbartolome, mica1014, micaelgabrielitliong, mjbirder, nathaniel_quinto, ninacam334, nopenot, ou_wildlife, oxfane, piper_andrea, popsie1, regina15366, rheinkayla_clemente, rossi107, salajumac, shea48, sophiedc, suebsa, sumimaxen, tonyg, timmyrii, tursiops, twan3253, wilhelmtan, wildjohnlee, yushe, zachary_gernes, zeb_amaro, zhedennar. Their collective efforts in documenting Lepidoptera and flowering plants across Metro Manila greatly enriched the dataset and strengthened the ecological insights presented in this work. This study stands as a testament to the power of citizen science in advancing urban biodiversity research. I would also like to extend my sincere thanks to Graham Joshua Ogatia for helping in the visualization of our datasets.

## Notes

### Competing Interest Statement

The authors have declared no competing interest.

